# Glutamine-Dependent Biosynthetic Pathways Fuel Autoreactive T and B Cells in Foxp3 Deficiency-mediated Disease

**DOI:** 10.1101/2025.11.04.686569

**Authors:** Mohammad Adeel Zafar, Charlotte Nicole Hill Machado, Jyotirmaya Behera, Yuelin Zhong, Xiao Li, Shakchhi Joshi, Yassine El Fazaa, Virginia Camacho, Peter Georgiev, Kiran Kurmi, Marcia Carmen Haigis, Louis-Marie Charbonnier

## Abstract

Foxp3 deficiency causes a profound loss of immune tolerance, unleashing autoreactive T and B cells, lymphoproliferation, cytokine-driven inflammation, and autoantibody production. This autoimmune pathology is fueled by increased glutamine usage, but it remains unresolved whether glutamine is necessary to produce energy, or for biosynthetic pathways leading to inosine and asparagine production. Here, we demonstrate that glutamine utilization supports Foxp3-deficiency mediated disease independently of pathogenic Foxp3-deficient Treg cell energetic reprogramming. Mechanistically, glutamine biosynthetic pathways sustain conventional T cell activation and proinflammatory cytokine production preventing inosine accumulation and signaling, thus implicating adenosine pathway modulation in autoreactive T cell dysregulation. Conversely, autoreactive B cell activation and autoantibody production depend on glutamine-dependent asparagine synthesis, which we reveal as a targetable vulnerability for autoantibody formation. These findings highlight glutamine-driven biosynthetic processes as critical drivers of autoimmunity and reveal distinct metabolic vulnerabilities in autoreactive T and B cells that can be targeted for therapeutic intervention.

## Introduction

Foxp3^+^ regulatory T (Treg) cells play a critical role in enforcing immune tolerance to self-antigens and benign antigens (1–3). Loss of function mutations in *FOXP3* in humans and its orthologue in mice result in a severe autoimmune lymphoproliferative disease, highlighting the crucial role of Foxp3 in Treg cell biology (4–7). Foxp3 deficiency does not impair T_reg_ cell development (8–10). While Foxp3-deficient Treg (ΔTreg) cells maintain a residual Treg signature, they also acquire effector T (Teff)-cell programs (6, 8, 9). The severity of Foxp3 deficiency mediated disease in mice can be reduced by CD4 or CD8 T cell depletion, Tbet deficiency, mature B cell deficiency, or Myd88 deficiency, implicating the contribution of innate immune cell maturation and activation and autoreactive T and B cells to disease pathogenesis (11–16).

Activation, differentiation and function of immune cells depends on their ability to adapt to and modify different cellular metabolic pathways due to their high energy demand. T and B cell activation requires metabolic reprogramming to meet the increased biosynthetic, bioenergetic, and signaling demands (17–20). Current autoimmune therapies largely focus on targeting inflammation or broadly inducing immunosuppression rather than addressing the underlying cause of disease (21–23). The activity of metabolic pathways is elevated in autoimmune diseases, and metabolic changes are increasingly recognized as important pathogenic processes underlying immune dysregulation. Therefore, metabolically targeted therapies may represent a conceptually novel strategy for treating autoimmune diseases (24). Key metabolic processes such as glycolysis, oxidative phosphorylation (OXPHOS), nucleotide and amino acid (AA) metabolism are vital for the immune cell survival and function. During inflammation, immune cells switch their metabolic status to meet the energy demand required for proliferation and effector molecule production (e.g. cytokines, Ab production, etc.). Several reports have highlighted a role of increased glutamine usage on immune cell activation in autoimmune diseases (25–30). In a previous study, we have shown that ΔTreg cells are highly proliferative and exhibit a Tconv-like metabolic profile associated with high glycolytic rate and high oxidative phosphorylation (OXPHOS). In contrast to Foxp3-sufficient Treg cells in which OXPHOS is primarily dependent on fatty acid oxidation (FAO), ΔTreg OXPHOS predominantly relies predominantly on glucose and glutamine to fuel mitochondrial energy production(9). These results bring glutamine metabolism to the forefront as one of the key metabolic targets for their therapeutic potential.

Glutamine, an abundant AA in the serum, serves as an essential fuel source for activated immune cells. Glutamine metabolism plays crucial roles in both bioenergetic and biosynthetic processes, particularly in rapidly dividing cells (31). Glutaminolysis fuels the bioenergetics of the cell by converting glutamine into glutamate by the enzyme glutaminase (GLS) and subsequently to α-ketoglutarate (αKG). αKG then enters the tricarboxylic acid (TCA) cycle, providing energy for cellular processes (32). Additionally, glutamine provides substrate for AA biogenesis and glutathione, required for protein synthesis and proliferation with the help of glutamic-oxaloacetic transaminase (GOT) or glutamate cysteine ligase (GCL), respectively (33, 34). Glutamine is required for de novo synthesis of purines and pyrimidines, which results from the activity of several enzymes including phosphoribosyl pyrophosphate amido transferase (PPAT) and carbamoyl-phosphate synthetase, aspartate transcarbamoylase, and dihydroorotase (CAD). Finally, glutamine also generates signaling molecules involved in activation of immune cells (35).

Here, we show that glutamine-dependent biosynthetic pathways drive autoreactive T and B cell responses in Foxp3-deficiency mediated disease. Surprisingly, systemic blockade of glutamine usage mitigates autoimmunity in a mouse model of Foxp3 deficiency not in a bioenergetic manner, but by altering glutamine-dependent biosynthetic pathways such as purine metabolites (inosine) and asparagine availability. Pharmacological inhibition of inosine signaling by A2A receptor and supplementation of asparagine specifically reverses the effect of glutamine blockade on T cells and B cells, respectively. Our work provides the basis for development of a new therapeutic approach to treat autoimmune and inflammatory diseases by inhibiting glutamine-dependent biosynthetic pathways.

## Results

### Glutamine-dependent biosynthetic pathways contribute to Foxp3 deficiency-mediated disease

To assess whether global glutamine utilization or only glutamine-dependent bioenergetic pathways were required to sustain autoimmunity, we initially compared the therapeutic effect of a broad glutamine antagonist 6-diazo-5-oxo-norleucine (DON) and of a glutaminase 1 (GLS1) specific inhibitor (CB-839) in the context of Foxp3 deficiency (*Foxp3*^ΔEGFPiCre^*R26*^YFP^ mice) by treating mice i.p every alternative day starting at D11 of age. Whereas DON broadly inhibits multiple glutamine-utilizing reactions essential for cellular processes like synthesis of nucleic acids and proteins, and generation of αKG for energy metabolism, CB-839 is a selective inhibitor of the enzyme glutaminase, converting glutamine into glutamate, thus preventing αKG production and utilization within the TCA cycle as well as glutamate-derived metabolites such as glutathione. We found that, DON-but not CB-839-treated mice showed a significant improvement in overall disease severity, body weight index and survival, as compared to vehicle (PBS) treated animals (**Figure 1A, B and Supplemental Figure 1A**). Cellular analysis revealed that DON, but not CB-839, significantly reduced the frequency and number of splenic ΔTreg cells (CD90^+^CD4^+^YFP^+^ cells), total CD4^+^ (CD90^+^CD4^+^CD8^−^ cells) and CD8^+^ (CD90^+^CD4^−^ CD8^+^ cells) T cell number, activated/memory (CD62L^lo^CD44^hi^) CD4^+^ and CD8^+^ Tconv (YFP^−^) cells frequency (**Figure 1C-F and Supplemental Figure 1B, C, F**). Furthermore, frequency and number of proinflammatory cytokine-producing CD4^+^ Tconv, ΔTreg (IFN-γ and IL-4) and CD8^+^ (IFN-γ) Tconv cells were also reduced upon DON treatment compared to vehicle (**Figure 1G, H and Supplemental Figure 1D-F**). Of note, Foxp3-deficient mice also have abundant autoantibodies (13, 36). We observed a significant reduction of splenic IgG1^+^IgD^−^ B cells, serum levels of total IgM, IgA, IgG1, IgG2b, IgG2c, IgG3 immunoglobulins (Ig) and direct IgG deposit within the kidney of DON-treated mice, as compared to vehicle-or CB-839-treated animals (**Figure 1I-M**). Indirect binding assays of serum IgG on ear sections of *Rag1*^KO^ mice revealed that only DON treatment, but not CB-839, reduced the autoreactivity of IgG on ear tissue, compared to serum from vehicle treated mice (**Supplemental Figure 1G,H**). To confirm the role of glutamine-dependent bioenergetic and biosynthetic pathways on B cell proliferation and class switching, we stimulated IgD^+^CD27^−^ naïve B cells isolated from untreated WT mice with LPS+IL-4, Resiquimod+IL-4, CD40L+IL-4 in presence of vehicle, CB-839 or DON *in vitro*. We observed that, while CB-839 reduced proliferation and class switching capacities of B cells *in vitro* in response to these 3 stimuli, *in vitro* DON exposure presented a superior inhibitory effect of B cell proliferation and Ab class switching, compared to vehicle (**Supplemental Figure 1I-P**). Furthermore, DON-treated *Foxp3*^ΔEGFPiCre^*R26*^YFP^ mice had decreased tissue inflammation (skin, liver, lungs and kidney), compared with vehicle-and CB-839-treated mice (**Figure 1N,O**). Together, these results show that overall blockade of glutamine functions, rather than its bioenergetic roles, support dysregulated autoreactive T and B cells to promote Foxp3 deficiency-mediated disease, suggesting that glutamine-dependent biosynthetic pathways play a predominant role in autoimmunity.

**Fig 1.**
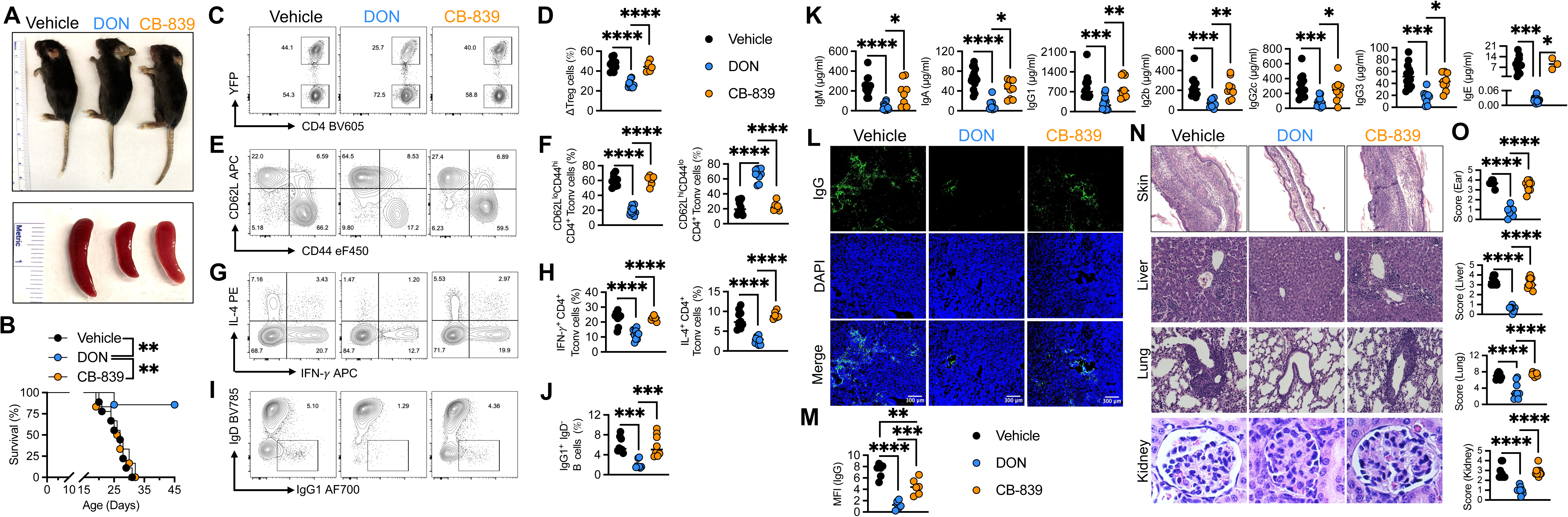
Glutamine-dependent biosynthetic pathways contribute to Foxp3 deficiency mediated disease. **A**,**B**, Gross appearance and their respective spleens of 21 days old *Foxp3*^ΔEGFPiCre^*R26*^YFP^ mice treated with vehicle, 6-diazo-5-oxo-norleucine (DON) or CB-839 (**A**), survival over time (**B**). The results represent one of the four experiments. **C-J,** Representative flow cytometric analysis and frequencies (scatter plots and means) of ΔTreg (CD90^+^CD4^+^CD8^−^YFP^+^) cells (**C,D**), CD62L^lo^CD44^hi^, CD62L^hi^CD44^lo^ CD4^+^ Tconv (CD90^+^CD4^+^CD8^−^YFP^−^) cells (**E,F**) and IFN-γ and IL-4 expression by CD4^+^ Tconv cells (**G,H**), IgG1^+^ IgD^−^B (CD19^+^) cells (**I,J**) from the spleen of mice of the respective treatment. **K**, Serum concentrations of immunoglobulin IgM, IgA, IgG1, IgG2b, IgG2c, IgG3 and IgE of mice of the respective treatment. **L**,**M**, Representative immunofluorescence pictures of IgG deposit (original magnification; ×200; (**L**) and IgG mean florescent intensity (MFI) (**M**) on kidney of mice of the respective treatment. (n=6 per group). **N,O**, Representative microscopic pictures of H&E staining (original magnification; ×200; (**N**) and histological scores (**O**) of the skin, lung, liver and kidney of mice of the respective treatment (n = 9). Statistical significance was determined by one-way ANOVA with Tukey’s multiple comparisons. *p < 0.05, **p < 0.01, ***p < 0.001, **** p < 0.0001.

### DON exerts beneficial effects independently of ΔTreg cells in Foxp3-deficient mice

Treg cells shift their metabolic status from fatty acid oxidation (FAO) to glycolysis and glutaminolysis upon loss of Foxp3, suggesting that ΔTreg cells at least partially rely on glutamine as a fuel for ATP production (9). Since *in vivo* glutaminolysis blockade by DON modulates T and B cell responses, to evaluate the contribution of ΔTreg cell reprogramming by DON treatment, we tested the impact of ΔTreg cell depletion in DON-treated Foxp3-deficient mice (**Figure 2**). To accomplish ΔTreg cell depletion, *Foxp3*^ΔEGFPiCre^*R26*^YFP^ and *R26*^iDTR^ mice were crossed and treated with DON alone or in combination with Diphtheria Toxin (DT). The effect of DON on gross appearance, splenomegaly, survival and body weight of Foxp3-deficient (*Foxp3*^ΔEGFPiCre^*R26*^YFP/iDTR^) mice was preserved even in association with DT treatment (**Figure 2A, B and Supplemental Figure 2A**). We validated that DT treatment i.p every alternative day from day 11 to day 21 in *Foxp3*^ΔEGFPiCre^*R26*^YFP/iDTR^ mice efficiently depleted the ΔTreg cell population (**Figure 2C, D**). Further, the effect of DON on CD4^+^ Tconv effector memory frequency and number, Th1/Th2 cytokine (IFN-γ/IL-4) production, CD4^+^ and CD8^+^ naïve T cell number, CD8^+^ Tconv effector memory frequency and IFN-γ production were marginally altered upon ΔTreg cell depletion with DT co-treatment (**Figure 2E-H; Supplemental Figure 2B-F**). A similar effect on IgG1 class switching of B cells, serum Ig levels, autoantibodies against skin antigens and IgG deposit in kidney by DON in presence or absence of ΔTreg cells was observed (**Figure 2K-M; Supplemental Figure 2G,H**). Finally, the overall effect of DON on tissue pathology was similar regardless of the presence of ΔTreg cells (**Figure 2N,O**). These results suggest that the salutary effects of DON on autoreactive and activated T and B cells are independent of ΔTreg cell metabolic reprogramming in the context of Foxp3 deficiency.

**Fig 2.**
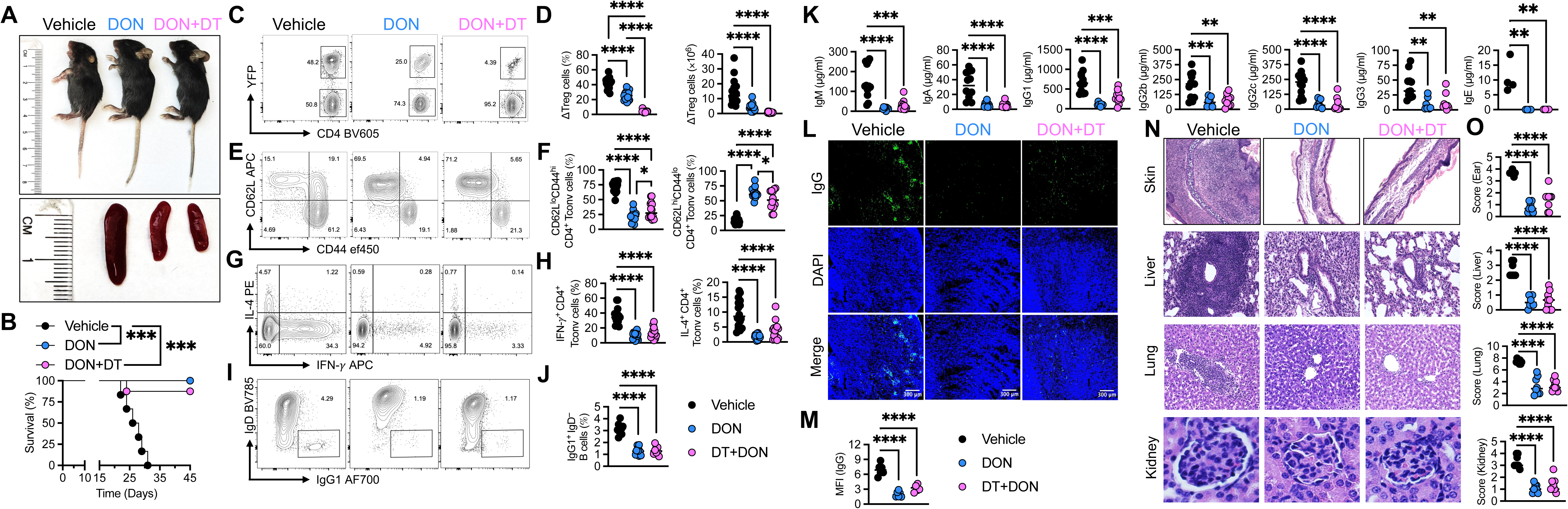
Glutamine usage by autoreactive T and B cell is essential for disease pathogenesis. **A**,**B**, Gross appearance and their respective spleens of 21 days old *Foxp3*^ΔEGFPiCre^*R26*^YFP/iDTR^ mice treated with vehicle, DON with or without diphtheria toxin (DT) (**A**), survival over time (**B**). The results represent one of the four experiments. **C-J,** Representative flow cytometric analysis and frequencies (scatter plots and means) of ΔTreg (CD90^+^CD4^+^CD8^−^YFP^+^) cells (**C,D**), CD62L^lo^CD44^hi^, CD62L^hi^CD44^lo^ CD4^+^ Tconv (CD90^+^CD4^+^CD8^−^YFP^−^) cells (**E,F**) and IFN-γ and IL-4 expression by CD4^+^ Tconv cells (**G,H**), IgG1^+^ IgD^−^B (CD19^+^) cells (**I,J**) from the spleen of mice of the respective treatment. **K**, Serum concentrations of immunoglobulin IgM, IgA, IgG1, IgG2b, IgG2c, IgG3 and IgE of mice of the respective treatment. **L**,**M**, Representative immunofluorescence pictures of IgG deposit (original magnification; ×200; (**L**) and IgG MFI (**M**) on kidney of mice of the respective treatment. (n=6 per group). **N,O**, Representative microscopic pictures of H&E staining (original magnification; ×200; (**N**) and histological scores (**O**) of the skin, lung, liver and kidney of mice of the respective treatment (n=9 per group). Statistical significance was determined by one-way ANOVA with Tukey’s multiple comparisons. *p < 0.05, **p < 0.01, ***p < 0.001, **** p < 0.0001.

### Glutamine usage blockage alters metabolic landscape

Various intermediate metabolites of glutamine metabolism contribute to different bioenergetic and biosynthetic processes. Glutamine acts as a source of carbon and nitrogen for nucleotide biosynthesis while its metabolite glutamate acts as a substrate for the synthesis of non-essential AAs (NEAAs) and Glutathione (37, 38). To further understand how inhibiting glutamine usage improves the Foxp3 deficiency-mediated disease, we analyzed *Foxp3*^ΔEGFPiCre^ mice’s plasma composition by LC-MS at day 15 after birth following vehicle or DON treatment on day 11 and 14. Using Principal Component Analysis (PCA), we were able to cluster the plasma samples by treatment (**Figure 3A**). Among 176 metabolites analyzed in vehicle-and DON-treated mouse plasma, 50 showed a significant difference in their relative abundance, represented in volcano plot and heatmap (**Figure 3B-D**). A pathway analysis (using Kyoto Encyclopedia of Genes and Genomes; KEGG) showed that the most altered metabolites were related to purine and histidine metabolism, suggesting that DON mostly affects these pathways (**Figure 3C,D**). Whereas metabolites related to the purine metabolism (e.g. FGAR, Guanosine, Inosine, Xanthine, Hypoxanthine,) were in higher relative abondance, metabolites related to glycolysis (e.g. Lactate, Pyruvate), TCA cycle (e.g. Citrate, Isocitrate) and glutamine-dependent AA synthesis (e.g. Alanine, Aspartate, Asparagine, Glutamate, Proline, Ornithine, Serine) were in lower relative abondance in DON-treated mice, compared to vehicle-treated ones (**Figure 3D**). Since DON, but not CB-839, mitigates Foxp3 deficiency-mediated disease, we concluded that the TCA cycle and other glutamine-dependent bioenergetic metabolic pathways do not play a critical role in the *in vivo* DON effect in Foxp3-deficient mice. Among the glutamine-dependent biosynthetic pathways, inosine and asparagine were significantly altered in differential abundance by DON treatment and represent interesting candidates to dampen respectively autoreactive T and B cell responses (39–42). Analysis of the top 25 compounds corelated with the inosine and asparagine suggested upregulated inosine metabolites and lower asparagine metabolites in the serum of DON-treated animals (**Figure 3E,F)**. These results support that glutamine generation of inosine and asparagine affects T and B cells, respectively.

**Fig 3.**
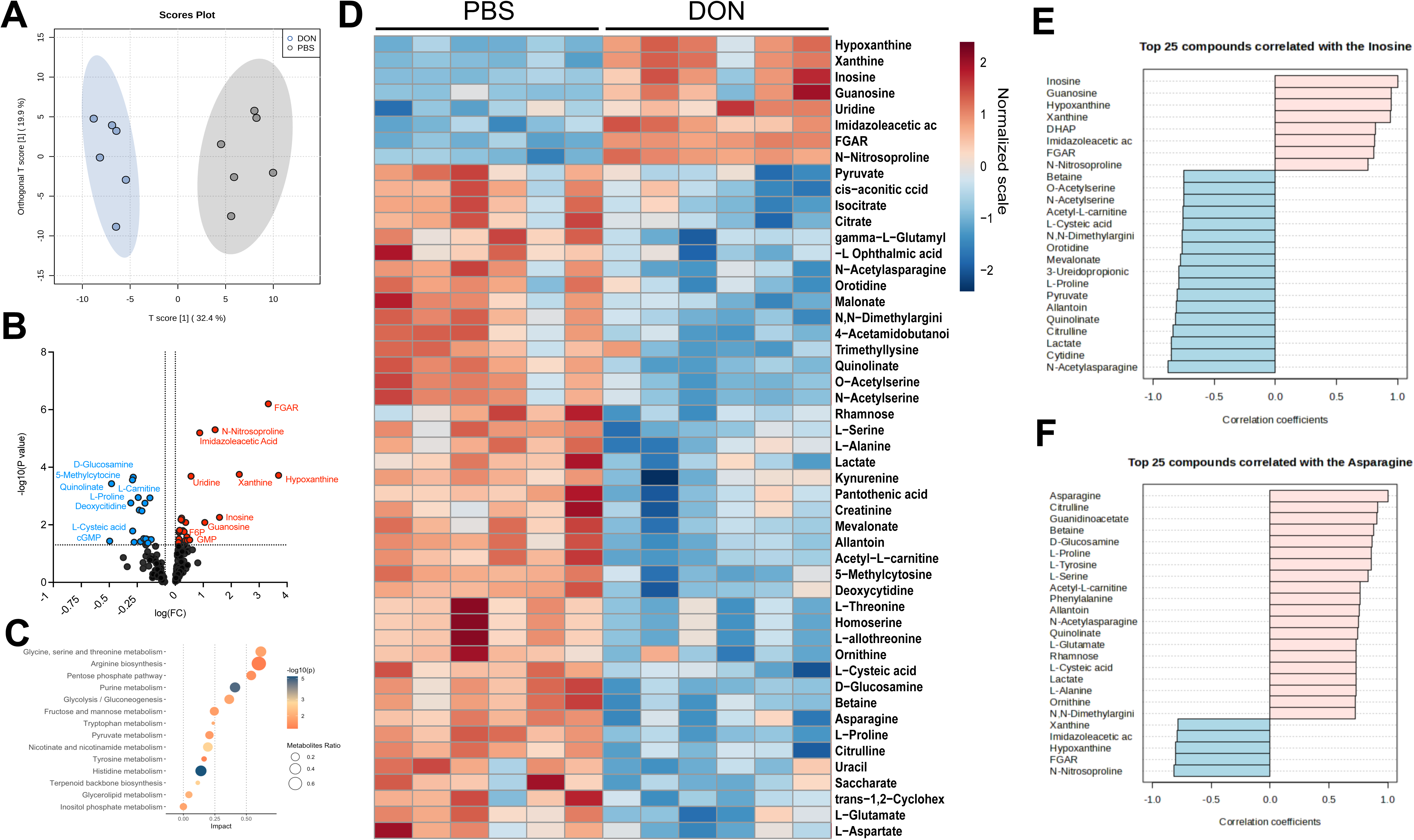
In-vivo Glutamine blockade alters purine and non-essential amino acid metabolism. **A,** PCA scores plot showing differentially expressed metabolites in DON vs PBS treated *Foxp3*^ΔEGFPicre^*R26*^YFP^ mice. **B**, loading plots showing metabolites upregulated (red) vs downregulated (blue) in the serum of DON treated *Foxp3*^ΔEGFPicre^*R26*^YFP^ mice. **C**, Lollipop diagram showing the significantly different KEGG pathways in DON vs PBS-treated mice plasma metabolites. **D**, heat map comparing the abondance of plasma metabolites in DON versus PBS treated Foxp3-deficient mice (DOWN and UP means that the metabolite abondance is respectively down or upregulated in the plasma of DON-treated mice, compared to PBS-treated ones). **E,F** Boxplots for pathway enrichment analysis showing the top 25 metabolites upregulated or downregulated related to inosine (**E**) and asparagine (**F**) metabolism in the serum of *Foxp3*^ΔEGFPicre^*R26*^YFP^ mice treated with DON.

### Inhibition of glutamine usage controls activation of autoreactive T and B cell in inosine and asparagine dependent manners respectively

Inosine binds the adenosine (A2A) receptor expressed on effector T cells to prevent Th1/Th2 skewing of T cells, while asparagine can be taken up directly by immune cells, particularly germinal center B cells (42–44). To confirm the role of altered inosine and asparagine levels on T and B cell activity following DON treatment, we treated *Foxp3*^ΔEGFPiCre^*R26*^YFP^ mice with DON in combination with an A2A receptor antagonist (SCH58261) or with asparagine supplementation. Co-treatment of DON with SCH58261 but not asparagine partially reversed the effect of DON on CD4^+^ Tconv effector memory frequency and number, Th1/Th2 cytokine (IFN-γ/IL-4) production by Tconv and ΔTreg cells, CD4^+^/CD8^+^/CD4^+^ effector T cell memory and naïve T cell number, CD8^+^ T cell number and IFN-γ production (**Figure 4A-F; Supplemental Figure 3A-H**). In contrast, while concurrent asparagine supplementation with DON marginally affected T cell responses, it reversed the effect of DON on B cell activation and autoreactivity, including restoration of IgG1 class switching of B cells, germinal center B cell formation, IgG/IgM autoreactivity against self-antigens, high serum Ig levels and IgG Ab deposit in kidney (**Fig. 4G-M; Supplemental Figure 3I**). As a result, concurrent A2A receptor blockade reversed the beneficial effect of DON on skin inflammation while upon asparagine supplementation restoration of IgG deposit in kidney resulted in increased severity of kidney pathology (**Figure 4N,O**). By comparing the *in vitro* capacities of naïve B cells isolated from untreated WT mice to proliferate and produce IgG1 upon LPS+IL-4 stimulation in presence or absence of DON and asparagine in asparagine low (DMEM) and rich (RPMI) medium, we confirm that asparagine is a critical NEAA for B cell proliferation and IgG1 production that can partially reverse the effect of DON (**Supplemental Figure 3J-L)**. Overall, these effects of A2A receptor blockade and asparagine supplementation in the presence of DON on T and B cells suggest that DON alters glutamine dependent biosynthetic pathways to inhibit T and B cell functions respectively via inosine and asparagine to mitigate the Foxp3 deficiency-mediated disease.

**Fig 4.**
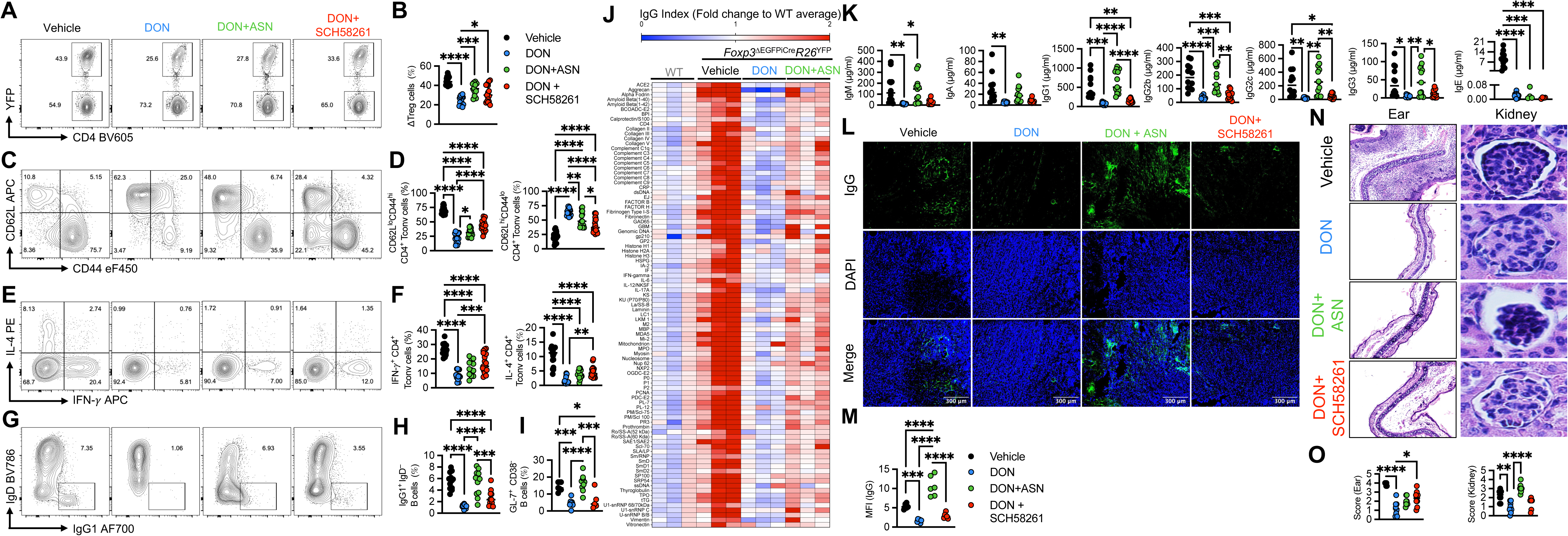
Inhibition of glutamine usage controls activation of autoreactive T and B cell in inosine and asparagine dependent manner respectively. **A-I,** Representative flow cytometric analysis and frequencies (scatter plots and means) of ΔTreg (CD90^+^CD4^+^CD8^−^YFP^+^) cells (**A,B**), CD62L^lo^CD44^hi^, CD62L^hi^CD44^lo^ CD4^+^ Tconv (CD90^+^CD4^+^CD8^−^YFP^−^) cells (**C,D**), IFN-γ and IL-4 expression by CD4^+^ Tconv cells (**E,F**), IgG1^+^ IgD^−^B (CD19^+^) cells (**G,H**), GL-7^+^ CD38^−^ germinal center B cells (**I**) from the spleen of mice treated with vehicle, DON, DON+ASN or DON+SCH58261. **J**, Heatmap of IgG reactivity to auto-antigen array analysis in the serum of untreated age-matched WT and *Foxp3*^ΔEGFPiCre^*R26*^YFP^ mice treated with vehicle, DON or DON+ASN. Fold change values for each group were calculated by normalizing with the values of WT mice. The observed fold change values are color-coded per the legend immediately to the top of the heatmap. **K**, Serum concentrations of immunoglobulin IgM, IgA, IgG1, IgG2b, IgG2c, IgG3 and IgE of mice of the respective treatment. **L**,**M**, Representative immunofluorescence pictures of IgG deposit (original magnification; ×200; (**L**) and mean florescent intensity (**M**) of kidney of mice of the respective treatment. (n = 6). **N,O**, Representative microscopic pictures of H&E staining (original magnification; ×200; (**N**) and histological scores (**O**) of the skin and kidney of mice of the respective treatment (n = 9). Statistical significance was determined by one-way ANOVA with Tukey’s multiple comparisons. *p < 0.05, **p < 0.01, ***p < 0.001, **** p < 0.0001.

## Discussion

Upon tolerance breakdown, immune cells proliferate intensively and produce inflammatory mediators (cytokines, chemokines, antibodies etc.). To exert these functions, they present disproportionate energy demand, and metabolic shift is the major adaptation to meet these demands. Recent understanding of metabolic reprogramming of immune cells in inflammatory conditions has paved the way for the development of novel therapeutics against inflammatory/autoimmune diseases. Many anti-inflammatory drugs such as metformin, rapamycin, dimethyl fumarate (DMF) and methotrexate can also directly impact immune cell metabolism by acting on different pathways including glucose metabolism, mTORC1, pentose phosphate pathway and fatty acid metabolism and nucleotide biosynthesis respectively (45). Interestingly, among different metabolic pathways, glutamine metabolism is still an underappreciated target for therapeutic control of immune cell activation. Breakdown of glutamine in proliferating cells serves as a carbon and nitrogen source to fuel bioenergetic and biosynthetic pathways. Deamination of glutamine to glutamate by glutaminase (GLS1/2) replenishes metabolic intermediate via TCA cycle for energy metabolism (46). Additionally, glutamine is involved in *de novo* synthesis of various metabolites such as nucleotides, NEAAs and glutathione (47). These molecules serve as a substrate for synthesis of effector molecules (cytokines, chemokines, antibodies etc.) required for T and B cell proliferation, survival and functions. Given the reliability of autoreactive T and B cells on glutamine-dependent biosynthetic pathways, these pathways can be targeted for treatment of inflammatory disorders.

In this study we used a Foxp3 deficiency model to study the contribution of dysregulated glutamine-dependent metabolic pathways used by autoreactive T and B cells to promote autoimmunity. Our results demonstrate that glutamine dependent biosynthetic but not bioenergetic pathways are essential to T and B cell activation/proliferation, inflammatory cytokines and autoantibody production in the context of Foxp3 deficiency. Inhibition of GLS1 by using CB-839 to target bioenergetic pathways has little to no effect on the severity of autoimmune disease mediated by Foxp3 deficiency, while targeting both glutamine-dependent bioenergetic and biosynthetic pathways using a broad inhibitor of glutamine utilizing enzyme DON resulted in overall improvement in the disease.

Plasma metabolite analysis revealed significant alteration in key metabolites (purine, pyrimidine, asparagine, etc.) related to glutamine metabolism in DON-treated mice suggesting the involvement of glutamine dependent biosynthetic pathways. Inhibition of glutamine-to-glutamate conversion resulted in higher purine and pyrimidine nucleotides, particularly inosine. Previous studies have demonstrated the anti-inflammatory effect of inosine as inosine treatment reduced the production of proinflammatory cytokines, chemokines and attenuated the course of chronic autoimmune inflammatory diseases including murine type I diabetes and experimental colitis (48, 49). Mechanistically, inosine binds the A2A receptor expressed on effector T cells to inhibit the Th1/Th2 polarization (40, 49, 50). Further, one study identified reduced levels of inosine in Foxp3-deficient mice, and found that replenishment of inosine resulted in improvement of the scurfy phenotype (40). We confirmed the role of elevated inosine levels on DON-mediated amelioration of disease by utilizing A2A receptor antagonist SCH58261. Co-administration of DON along with SCH58261 resulted in partial reversal of DON-mediated effects on T cells.

Glutamate also acts as a nitrogen donor for the synthesis of NEAA. Inhibition of glutamine metabolism by DON resulted in reduced NEAA abundance. Among various NEAAs that were significantly reduced in the plasma of DON treated animals, asparagine was the most promising candidate for its role in the B cell activation and germinal center formation (42). Our results indicated asparagine as key regulator of B cell homeostasis specially in autoAb production. In this study, we showed that glutamine-dependent asparagine availability in Foxp3 deficient mice is critical for B cell autoreactivity and autoAb production, and by targeting glutamine-dependent asparagine synthesis, we reduced the B cell autoreactivity in the context of Foxp3 deficiency, suggesting a critical role of asparagine in B cell dysregulated responses in autoimmune settings.

DON has been studied for decades as a potential anti-cancer therapeutic. Our result showed that it can be useful for the treatment of inflammatory disorders and autoimmunity largely due to its effect on glutamine dependent biosynthetic pathways responsible for generation of metabolic intermediates required for activation of T and B cells. More broadly, our studies uncover the possibility of combinatorial interventions that target distinct metabolic pathways to restore immune tolerance in a variety of autoimmune and immune dysregulatory diseases.

## Methods

### Animals and treatment

C57BL/6 WT and *Rag1*^KO^ were purchased to Jackson Laboratory. *Foxp3*^ΔEGFPiCre^*R26*^YFP^ strain was generated as described previously (9)*, Foxp3*^ΔEGFPiCre^*R26*^YFP/iDTR^ mice were generated by crossing *Foxp3*^ΔEGFPiCre^*R26*^YFP^ with *R26*^iDTR^ (Jackson Laboratory). Mice were treated with DON, CB-839, SCH58261 and asparagine (3μg/g, 12.5μg/g, 6μg/g and 24μg/g of bodyweight respectively), intraperitoneally for every alternate day from day 11 to 21. *Foxp3*^ΔEGFPiCre^*R26*^YFP/iDTR^ mice were treated with diphtheria Toxin (DT) (0.5mg/mouse on day 11 and 0.25mg/mouse from day 12 onwards) intraperitoneally for every alternate day up to day 21. Animals were sacrificed at D22 and serum, spleen, ear, lung, liver and kidney were collected for autoantibody array, antibody ELISA, flowcytometry, histopathology and immunohistochemistry. Animals were kept in specific pathogen free condition for the entire duration of experiment and used in accordance with institutional animal research committees at the Boston Children’s Hospital.

### Reagents

6-Diazo-5-oxo-L-norleucine (DON) (#Catalog D2141), Diphtheria Toxin (#Catalog D0564), L-Asparagine (#Catalog A0884) were acquired from Sigma Aldrich. SCH58261 was purchased from Tocris (#Catalog 2270). CB-839 was purchased from Cayman Chemical Company (#Catalog 22038). All the reagents were diluted and stored according to manufacturer’s instruction.

### B cell class switching and proliferation assay

IgD^+^CD27^−^ naïve B cells were purified from WT mice by using Dynabeads™ Mouse CD43 kit (invitrogen) according to manufacturer’s instruction. Cell Trace violet labelled 5×10^4^ naïve B cells were cultured in RPMI medium supplemented with 10% FBS, 1mM sodium pyruvate,10mM HEPES, 100 IU/ml penicillin/streptomycin and 50μM 2-mercaptoethanol, Aspartate (150μM), Glutamate (140μM), GlutaMAX (2mM) for 48 hours in presence or absence of LPS (10μg/ml) / Resiquimod (5μg/ml) / CD40L (5μg/ml) with IL-4 (50ng/ml) in 96 well plate. DMEM without NEAA was used as ASN free medium wherever indicated. Cells were treated with either DON/ASN or in combination as indicated in the figures. For proliferation assay, after Fc block, cells were stained with viability dye and CD19. For class switching assay, splenocytes or cultured cells were incubated with monoclonal anti IgE antibody along with Fc block followed by staining with viability dye and surface markers against CD19, IgD and later stained intra cellularly with anti-IgG1, IgM and IgE antibodies.

### Flow cytometry

Cells were stained as previously described(9). Various anti-mouse antibodies against CD44 (eBioscience), CD90.1, CD4, CD8a, CD62L, IFN-γ, IL-4, CD19, IgD, IgG1, IgM along with viability dye and Fc block (CD16/32) (Biolegend) were used. Briefly, cells were incubated for 10 min with Fc Block followed by viability dye and surface markers for 20 min. For intracellular markers, after surface staining, cells were fixed and permeabilized with BD Cytofix/Cytoperm buffer for 30 min and incubated overnight with antibodies against intracellular markers. For cytokine detection cells were pre-stimulated with 50 ng/ml phorbol myristate acetate (PMA), 500 ng/ml ionomycin and 10 μg/ml brefeldin A for 4 h at 37°C in RPMI medium with 10% FBS. After staining cells were acquired using BD Fortessa with DIVA software (BD Biosciences) and analyzed using FlowJo software. All the antibodies used for flowcytometry are listed in the **supplementary table 1**.

### Histopathology

Ear, kidney, lung and liver sections were stained with Hematoxylin and Eosin. Bright field Images at 200× magnification were taken on EVOS M700 (Invitrogen) microscope. Histological scores were performed by taking an average of 3 different fields of each section by a blinded observer. Ear inflammation was scored as follows: 0: no inflammation; 1: mild inflammation associated with infiltration of a few cells; 2: moderate inflammation associated with mild infiltration; 3: severe inflammation associated with large infiltration of cells and mild skin dryness; and 4: very severe inflammation associated with skin dryness and cartilage erosion. The glomerular lesions were graded on a scale of 0– 3 as previously described (51). Histopathologic findings in renal vascular lesions were graded on a scale of 0–3: 0: normal; 1: mild (perivascular cell infiltration); 2: moderate (destruction of arterial wall); 3: severe (myointimal thickening). Lung inflammation was scored separately for cellular infiltration around blood vessels and airways, as follows: 0: no infiltrates; 1: few inflammatory cells; 2: a ring of inflammatory cells one cell layer deep; 3: a ring of inflammatory cells two to four cells deep; and 4: a ring of inflammatory cells more than four cells deep. A composite score was determined by adding the inflammatory scores for both vessels and airways. Liver inflammation was scored at portal areas, as follows: 0: no inflammatory cells; 1: mild, scattered infiltrates; 2: moderate infiltrates occupying <50% of the portal areas; 3: extensive infiltrates in the portal areas; and 4: severe, with infiltrates completely packing the portal area and spilling over into the parenchyma.

### Immunohistochemistry

Kidney cryosections were fixed with acetone followed by blocking with 2% BSA and stained with fluorochrome conjugated anti-mouse IgG Ab for evaluating direct Ab deposit in kidney. Ear cryosections from *Rag1*^KO^ mice were used to detect autoreactive antibodies against skin antigens. After fixation and blocking, ear samples were incubated with serum (1:30) from 22 days old mice of the respective genotypes and treatments for 1 hour followed by incubation with anti-mouse IgG Ab for 30 minutes. Finally, sections were mounted using VECTASHIELD PLUS antifade mounting medium with DAPI (Vector Laboratories) and images were collected using fluorescence microscope and processed using ImageJ software. Final scores reflected averages of scores from three different ×200 fields per tissue per mouse.

### Antibody ELISA

ELISA was performed to detect total IgE/IgG1 antibodies in the culture supernatant as described (52). ELISA plates coated overnight with capture antibodies were blocked with 2% BSA solution. Supernatant samples and standards were incubated at room temperature for 2 hours. Next, biotin conjugated secondary antibodies were added for 1 hour followed by incubation with avidin HRP solution (Biolegend) for 45 minutes. Finally, wells were developed by using TMB substrate (BD Biosciences) and the reaction was stopped using 1M HCl and read at 450nm using spectrophotometer.

### Mouse immunoglobulin isotyping panel detection

Concentration of IgG1, IgG2b, IgG2c, IgG3, IgA, IgM, IgE Ig isotypes in the serum of different group was measured using bead based LEGENDplex™ Mouse Immunoglobulin Isotyping Panel (Biolegend) according to manufacturer instruction.

### Metabolomics analysis by LC-MS

*Foxp3*^ΔEGFPiCre^*R26*^YFP^ mice received intraperitoneal injections of DON (0.3mg/kg), or PBS on day 11 and 14. On day 15, whole blood was collected in K2-EDTA tubes and centrifuged at 1,500 x gravity for 15 min at 4°C. Metabolite extraction was then performed on the plasma fraction by adding 1mL of −20°C methanol and chloroform solvent (60:40) and vortexed for 15 minutes at 4°C and centrifuged at 21,000 x gravity for 15 min at 4°C. The supernatant containing polar metabolites was dried using a 4°C CentriVap SpeedVac (Labconco). Dried metabolite samples were resuspended in 50:50 acetonitrile:water for LC-MS analysis. Metabolites were resolved on a Vanquish U-HPLC system coupled to a Q Exactive HF-X hybrid quadrupole-orbitrap mass spectrometer (ThermoFisher) with a HESI source operating in negative ion mode. The analytes were separated by using an iHILIC column (5mm, 150 3 2.1 mm I.D., HILICON) coupled to a Thermo Scientific SII UPLC system. The iHILIC column was used with the following buffers: A = water with 20 mM ammonium carbonate with 0.1% ammonium hydroxide, B = acetonitrile. The Vanquish U-HPLC was run at a flow rate of 0.150 mL/min: 0-23 min linear gradient from 95% B to 5% B; 23-25 min hold at 5% B, to waste from 25-25.5 min gradient to 95% B at 0.20 mL/min, 25.5-32.5 min hold at 95% B, and finally 32.5-33 min 95% B at 0.15 mL/min. MS data targeted feature extraction and quantification were performed on TraceFinder v4.1 (ThermoFisher). Peak area integration and metabolite identification were performed using accurate mass and retention time curated with in-house standard library compounds. Data were normalized by cell number, and internal standard D8-phenylalanine was spiked during metabolite extraction to account for variations introduced during sample handling, preparation, and injection. MetaboAnalyst 5.0 platform was used for metabolomics data analysis.

### Autoantibody array

AutoAb autoantigen (anti-mouse IgG, and anti-mouse IgM) screening against 90 autoantigens in the serum samples was performed by Core Facility of the University of Texas Southwestern Medical Center (UTSW; Dallas, Texas, USA) as previously described (53).

## Acknowledgments

This work was supported by NIH NIAID grant R01AI153174 (to L-M.C).

## Authorship

### Contribution

M.A.Z., M.C.H and L.-M.C. designed experiments. M.A.Z., C.N.H.M., J.B, Y.Z., X.L., S.J., Y.E.F., V.C., P.G., K.K., and L.-M.C. performed experiments. P.G. and K.K. analyzed metabolomic results. M.A.Z and L.-M.C. made the figures and wrote the manuscript.

---

**Supplemental Figure 1. A.** Body weight change of the *Foxp3*^ΔEGFPiCre^*R26*^YFP^ mice treated with vehicle, DON and CB-839 over time. **B-E,** Representative flow cytometric analysis and frequencies (scatter plots with means) of CD62L^lo^CD44^hi^ and CD62L^hi^CD44^hi^ CD8^+^ Tconv (CD90^+^CD4^−^CD8^+^YFP^−^) cells (**B,C**), IFN-γ expression by CD8^+^ Tconv cells (**D,E**) from the spleen of the respective treatment group. **F**, Absolute count of total CD4^+^ (CD90^+^CD4^+^CD8^−^) and CD8^+^ (CD90^+^CD4^−^CD8^+^) T cells, ΔTreg (CD90^+^CD4^+^CD8^−^YFP^+^) cells, CD62L^lo^CD44^hi^, CD62L^hi^CD44^lo^ CD4^+^ Tconv (CD90^+^CD4^+^CD8^−^YFP^−^) cells and frequency of IFN-γ and IL-4 expression by ΔTreg cells. **G,H** Representative indirect immunofluorescence (**G**) and quantification (IgG MFI) (**H**) of the incidence of circulating tissue-specific antibodies (IgG) in the serum of *Foxp3*^ΔEGFPiCre^*R26*^YFP^ mice treated with DON, CB-839 or vehicle, using *Rag1*^KO^ mouse ear cryosections (n=6 per group). **I-P**, Representative histogram (**I**) frequencies (**J**) of CellTrace Violet^lo^ (divided) B (CD19^+^) cells, representative flowcytometric plot (**K, N**) and frequencies (scatter plots with means) (**L, O**) of B cell class switching (total IgM, IgE, IgG1) and Ig isotype levels in supernatant (total IgG1 and IgE) (**M,P**) of B cells stimulated with LPS, Resiquimod, CD40L+IL-4 cultured for 48h with vehicle, DON or CB-839. Statistical significance was determined by one-way ANOVA with Tukey’s multiple comparisons. *p < 0.05, **p < 0.01, ***p < 0.001, **** p < 0.0001.

**Supplemental Figure 2. A.** Body weight change of *Foxp3*^ΔEGFPiCre^*R26*^YFP/iDTR^ mice treated with vehicle, DON with or without diphtheria toxin (DT) over time. **B-E,** Representative flow cytometric analysis and frequencies (scatter plots with means) of CD62L^lo^CD44^hi^ CD8^+^ Tconv (CD90^+^CD4^−^CD8^+^YFP^−^) cells (**B,C**), IFN-γ expression by CD8^+^ Tconv cells (**D,E**) from the spleen of the respective treatment group. **F**, Absolute count of total CD4^+^ (CD90^+^CD4^+^CD8^−^) and CD8^+^ (CD90^+^CD4^−^CD8^+^) T cells, CD62L^lo^CD44^hi^, CD62L^hi^CD44^lo^ CD4^+^ Tconv (CD90^+^CD4^+^CD8^−^YFP^−^) cells and frequency of IFN-γ and IL-4 expression by ΔTreg (CD90^+^CD4^+^CD8^−^YFP^+^) cells. **G,H** Representative indirect immunofluorescence (**G**) and quantification (mean fluorescence intensity, MFI) (**H**) of the incidence of circulating tissue-specific antibodies (IgG) in the serum of *Foxp3*^ΔEGFPiCre^*R26*^YFP/iDTR^ mice treated with vehicle, DON with or without diphtheria toxin (DT), using *Rag1*^KO^ mouse ear cryosections (n=6 per group). Statistical significance was determined by one-way ANOVA with Tukey’s multiple comparisons. *p < 0.05, **p < 0.01, ***p < 0.001, **** p < 0.0001.

**Supplemental Figure 3. A.** Gross appearance and their respective spleens of 21 days old *Foxp3*^ΔEGFPiCre^*R26*^YFP^ mice treated with vehicle, DON, DON+asparagine (ASN) or DON+SCH58261. **B-G,** Representative flow cytometric analysis and frequencies (scatter plots with means) of CD62L^lo^CD44^hi^ CD8^+^ Tconv (CD90^+^CD4^−^CD8^+^YFP^−^) cells (**B,C**), IFN-γ expression by CD8^+^ Tconv cells (**D,E**), IFN-γ and IL-4 expression by ΔTreg (CD90^+^CD4^+^CD8^−^YFP^+^) cells (**F,G**) from the spleen of the respective treatment group. **H**, Absolute count of total CD4^+^ (CD90^+^CD4^+^CD8^−^) and CD8^+^ (CD90^+^CD4^−^CD8^+^) T cells, ΔTreg cells, CD62L^lo^CD44^hi^, CD62L^hi^CD44^lo^ CD4^+^ Tconv (CD90^+^CD4^+^CD8^−^YFP^−^) cells from the spleen of the respective treatment group. **I**, Heatmap of IgM reactivity to auto-antigen array analysis in the serum of untreated age-matched WT and *Foxp3*^ΔEGFPiCre^*R26*^YFP^ mice treated with vehicle, DON or DON+ASN. **J-L**, Representative histogram (**J**) frequencies (**K**) of CellTrace Violet^lo^ (divided) B (CD19^+^) cells and IgG1 level in the supernatant (**L**) of B cell cultured in ASN rich (RPMI) or ASN low (DMEM) medium treated with vehicle, DON, ASN or DON+ASN. Statistical significance was determined by one-way ANOVA with Tukey’s multiple comparisons. *p < 0.05, **p < 0.01, ***p < 0.001, **** p < 0.0001.

**Supplementary Table 1.** Listing of antibodies used in the study.

| **Marker** | **Fluorochrome** | **clone** | **company** | **dilution** | **staining** |
| --- | --- | --- | --- | --- | --- |
| Viability | eFluor506 |  | eBiosciences | 1:1000 | Surface |
| CD90.2 | APC-Cy7 | 30-H12 | Biolegend | 1:500 | Surface |
| CD4 | BV605 | GK1.5 | Biolegend | 1:500 | Surface |
| CD44 | eF450 | IM7 | eBiosciences | 1:500 | Surface |
| CD62L | APC | MEL-14 | Biolegend | 1:500 | Surface |
| IFN-γ | APC | XMG1.2 | Biolegend | 1:500 | Intracellular |
| IL-4 | PE | 11B11 | Biolegend | 1:500 | Intracellular |
| CD8 | Percp-cy5.5 | 53-6.7 | Biolegend | 1:500 | Surface |
| CD19 | APC-Fire750 | 6D5 | Biolegend | 1:500 | Surface |
| IgD | BV785 | 11-26c.2a | Biolegend | 1:500 | Surface |
| CD27 | PE | LG.3A10 | Biolegend | 1:500 | Surface |
| IgG1 | AF700 | RMG1-1 | Biolegend | 1:500 | Intracellular |
| IgE | BV421 | R35-72 | BD Biosciences | 1:500 | Intracellular |
| IgM | BV605 | RMM-1 | Biolegend | 1:500 | Intracellular |
| CD38 | PE-Cy7 | S21016F | Biolegend | 1:500 | Surface |
| GL-7 | Percp-cy5.5 | GL-7 | Biolegend | 1:500 | Surface |
| CD16/32 | Unconjugated | S17011E | Biolegend | 1:200 | Surface |
| IgE | Unconjugated | R35-72 | BD Biosciences | 1:300 | Surface |

