## Supplementary figures and images for "Glutamine-Dependent Biosynthetic Pathways Fuel Autoreactive T and B Cells in Foxp3 Deficiency-mediated Disease"

### Supplementary Figures 1 to 3

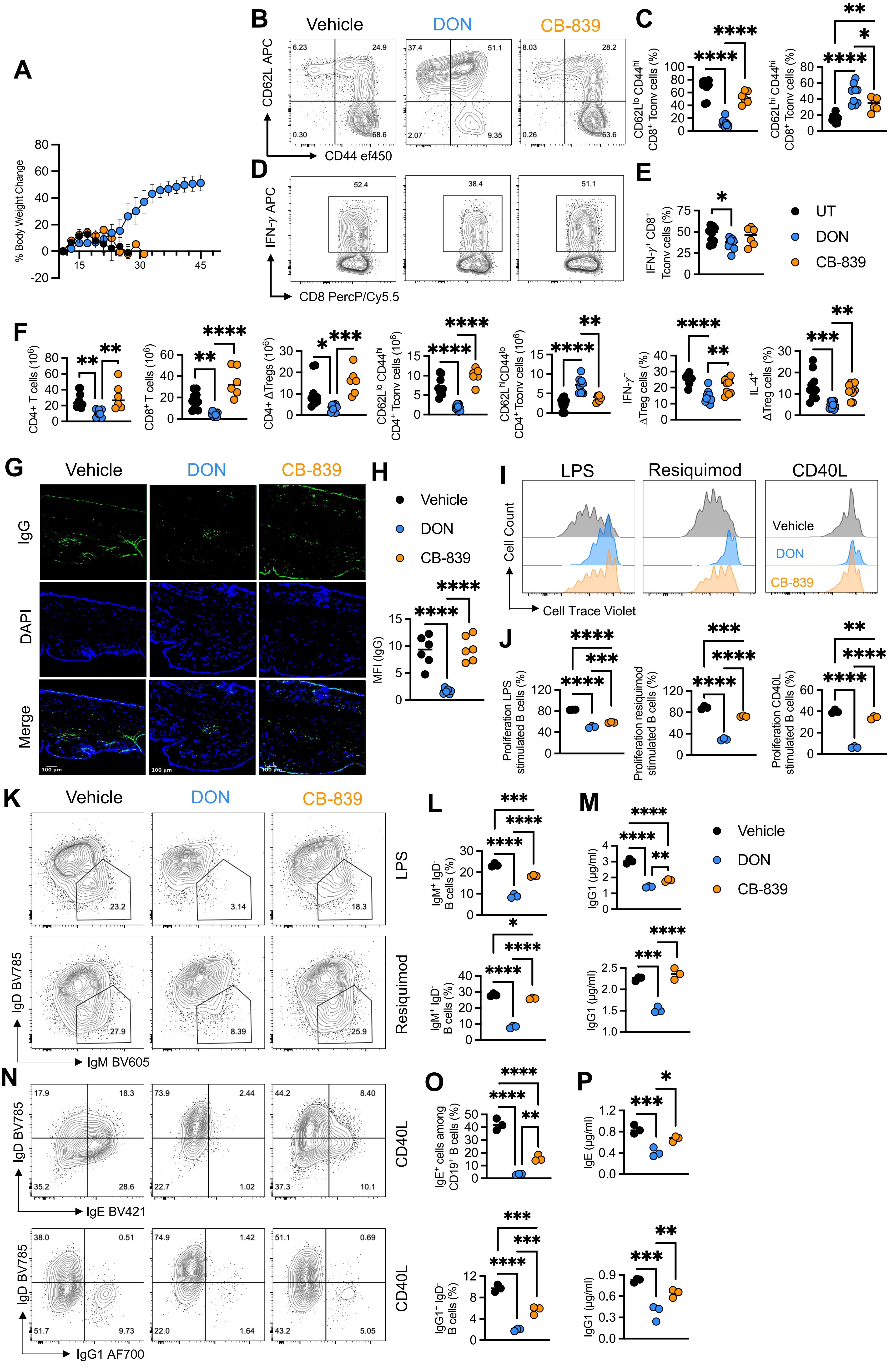

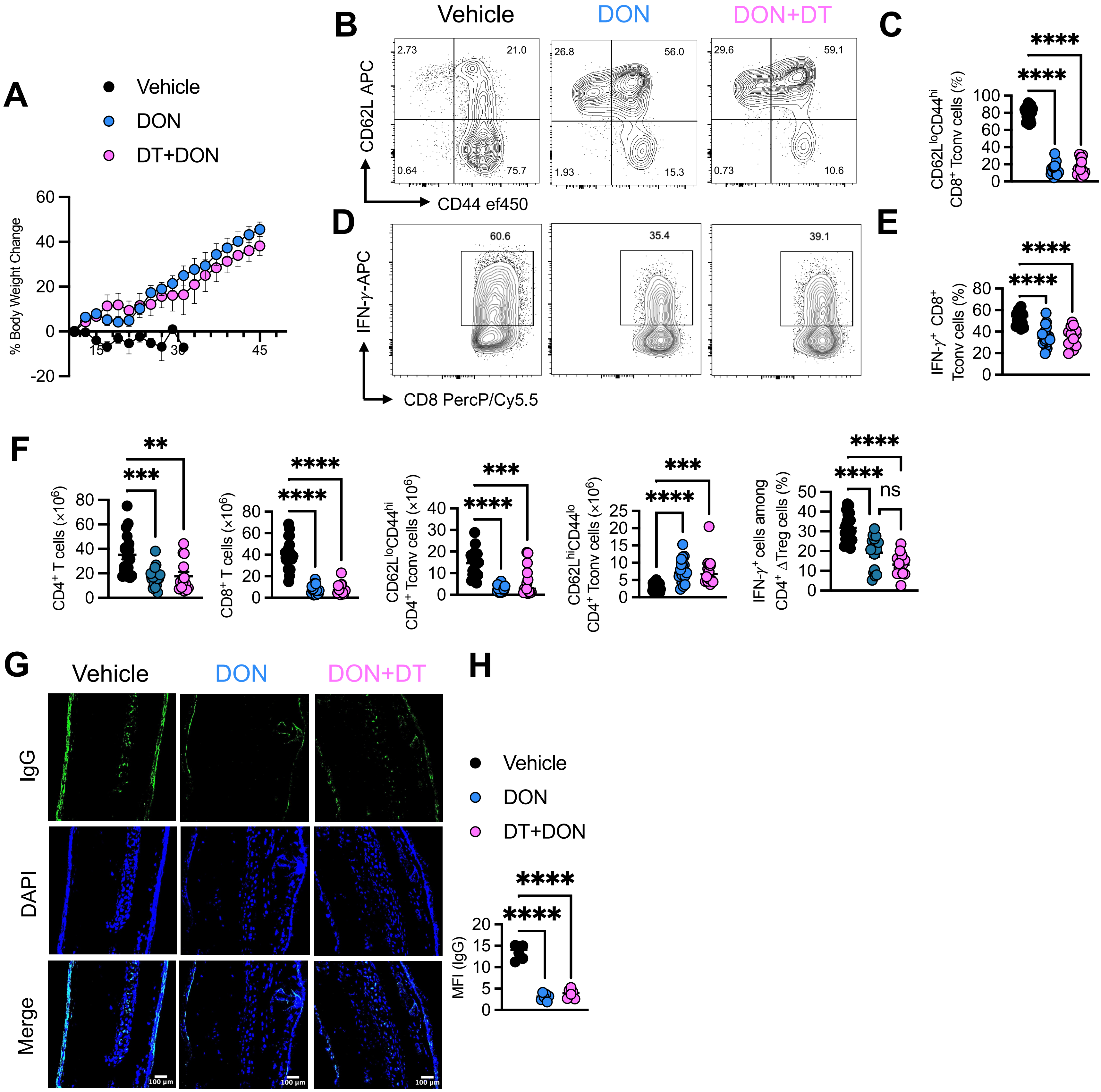

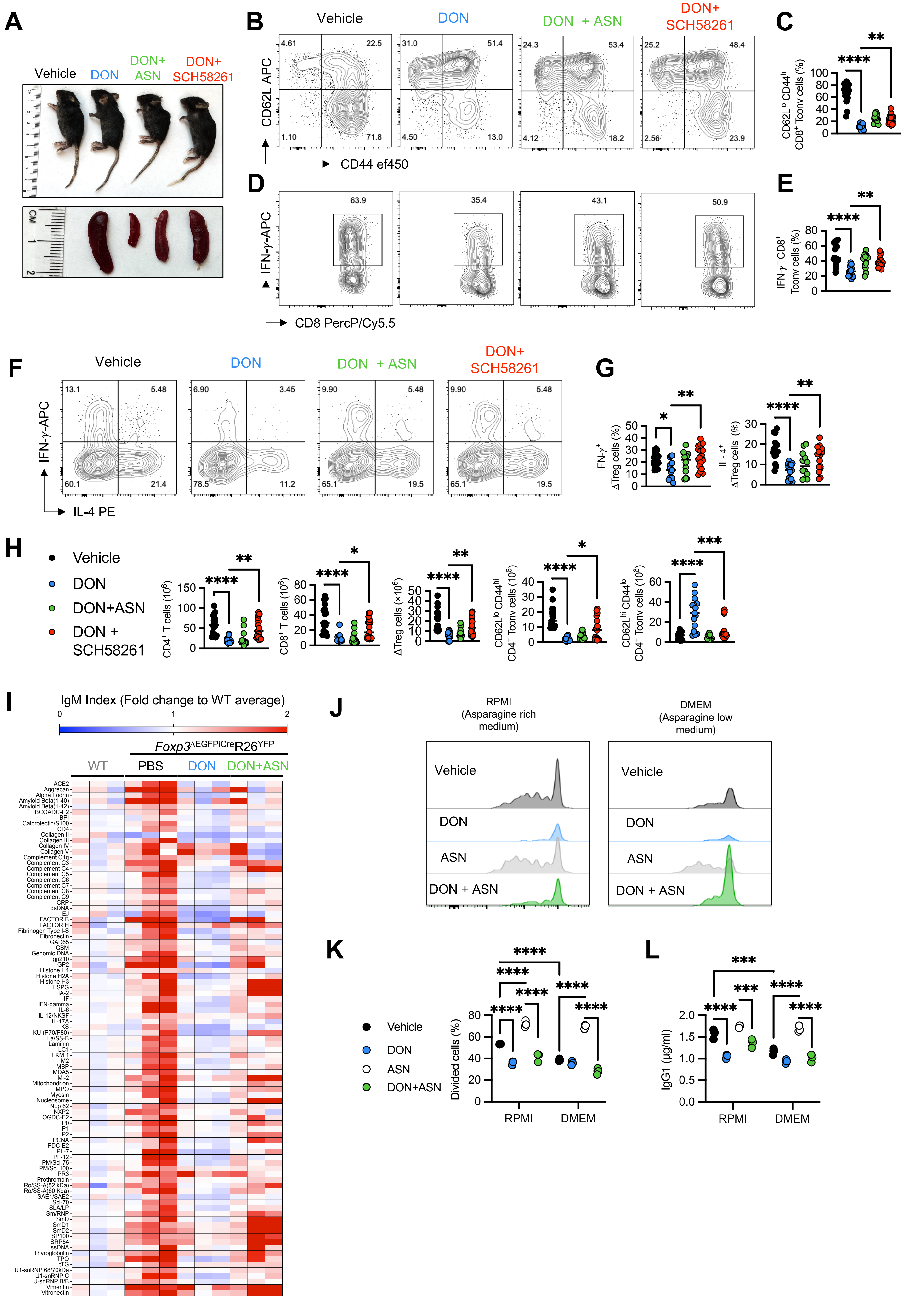
